# Distinct genetic bases for plant root responses to lipo-chitooligosaccharide signal molecules from distinct microbial origins

**DOI:** 10.1101/2020.09.09.285668

**Authors:** Maxime Bonhomme, Sandra Bensmihen, Olivier André, Emilie Amblard, Magali Garcia, Fabienne Maillet, Virginie Puech-Pagès, Clare Gough, Sébastien Fort, Sylvain Cottaz, Guillaume Bécard, Christophe Jacquet

## Abstract

- Lipo-chitooligosaccharides (LCOs) were originally found as symbiotic signals called Nod Factors (Nod-LCOs) controlling nodulation of legumes by rhizobia. More recently LCOs were also found in symbiotic fungi and, more surprisingly, very widely in the kingdom fungi including in saprophytic and pathogenic fungi. The LCO-V(C18:1, Fuc/MeFuc), hereafter called Fung-LCOs, are the LCO structures most commonly found in fungi. This raises the question of how legume plants, such as *Medicago truncatula*, can perceive and discriminate between Nod-LCOs and these Fung-LCOs.
- To address this question, we performed a Genome Wide Association Study on 173 natural accessions of *Medicago truncatula*, using a root branching phenotype and a newly developed local score approach.
- Both Nod- and Fung-LCOs stimulated root branching in most accessions but there was very little correlation in the ability to respond to these types of LCO molecules. Moreover, heritability of root response was higher for Nod-LCOs than for Fung-LCOs. We identified 123 loci for Nod-LCO and 71 for Fung-LCO responses, but only one was common.
- This suggests that Nod- and Fung-LCOs both control root branching but use different molecular mechanisms. The tighter genetic constraint of the root response to Fung-LCOs possibly reflects the ancestral origin of the biological activity of these molecules.

## Introduction

Lipo-chitooligosaccharides (LCOs) belong to a family of chitin oligomers substituted on their non-reducing end with an acyl chain, and further substituted with a variety of additional functional groups. LCOs were originally found, 30 years ago, to be symbiotic signals, called Nod factors, produced by rhizobia to trigger the nodulation process in legumes (Dénarié *et al*., 1996). This discovery was the starting point for a series of work that gradually brought to light the symbiotic signaling pathway required for rhizobial infection and nodulation in legumes. The activation of this signaling pathway, now called the Common Symbiosis Signalling Pathway (CSSP), was also found to be necessary for root colonization by arbuscular mycorrhizal (AM) fungi (Catoira *et al*., 2000). Furthermore, it was subsequently discovered that LCOs with high structural similarity to Nod factors are also produced by AM fungi (so called Myc-LCOs, Fig. S1) (Maillet *et al*., 2011). Without genetic proof that these molecules are essential for mycorrhization, but since they activate the CSSP as well as symbiotic gene expression changes in host plants, they are considered, together with their oligosaccharidic precursors (COs), as key mycorrhizal signals (Gough & Cullimore, 2011; Genre *et al*., 2013; Camps *et al*., 2015; Sun *et al*., 2015). This is supported by the recent finding in *Solanum lycopersicum*, that the receptor protein SlLYK10 binds Myc-LCOs and controls the AM symbiosis (Girardin *et al*., 2019). Also recently, Cope et al. showed both that the CSSP is used for establishment of the ectomycorrhizal symbiosis between *Laccaria bicolor* and poplar, and that *L. bicolor* can produce LCOs with similar structures to Nod factors (Cope *et al*., 2019). Possibly linked to their roles as symbiotic signals, LCOs can interfere with immunity-related signaling in legumes (Rey *et al*., 2019) and suppress innate immune responses, even in the non-mycorrhizal plant *Arabidopsis thaliana* (Liang *et al*., 2013). How LCOs dampen legume immunity is still unclear and controversial since they can also induce defense gene expression (Nakagawa *et al*., 2011). Another property of LCOs is their ability to modify root architecture by stimulating Lateral Root Formation (LRF). The stimulation of LRF appears to be a general response, observed in legume species such as *Medicago truncatula* treated with Nod Factors or Myc-LCOs (Olah *et al*., 2005; Maillet *et al*., 2011), but also in the monocots rice and *Brachypodium distachyon* (Sun *et al*., 2015; Buendia *et al*., 2019). Other positive effects of LCOs on soybean or maize root development are reported (Souleimanov *et al*., 2002; Tanaka *et al*., 2015). So, up to this point in our knowledge, LCOs were considered as signal molecules produced by a variety of symbiotic microorganisms and with several effects on plants, including activation of the CSSP, regulation of immune responses and stimulation of root development.

However, very recently, a new LCO chapter was opened when Rush et al. (Rush *et al*., 2020) discovered both that AM fungi produce a wider range of LCOs than previously described, and that LCOs are not exclusive to symbiotic microorganisms, but are actually a family of molecules commonly produced by a very large number of fungi, in all clades of the fungi kingdom. As such, they will be thereafter referred to as “Fung-LCOs”. Like previously characterized LCOs, Fung-LCOs consist of oligomers of 3-5 residues of *N*-acetyl glucosamine acylated with fatty acid chains of various length, saturated or not, and are decorated with acetyl, *N* methyl, carbamoyl, fucosyl, fucosyl sulfate, methyl fucosyl or sulfate groups. They can be found in phytopathogenic fungi, but also in saprophytes and opportunistic human pathogens, *i*.*e*. in non-symbiotic fungi or in fungi that do not interact with plants. The results of Rush et al. suggest that Fung-LCOs are conserved molecules in fungi that can regulate endogenous developmental processes such as spore germination, hyphal branching, or dimorphic switching. The fact that LCO-producing fungi of all kinds are abundantly present in the close environment of plant roots raises many new questions.

Focusing on the plant side, some of these questions might be: are these Fung-LCO structures able to trigger similar root responses, especially the LRF stimulation previously observed in response to Nod- and Myc-LCOs? If so, are legumes nevertheless able to differentiate these Fung-LCOs from the Nod-LCOs? To address these questions, we used a natural variability approach to compare root growth responses to Fung-LCOs and Nod-LCOs, using the model plant *Medicago truncatula*. As a legume, this plant must distinguish between Nod factors specifically produced by its rhizobial symbiont, *Sinorhizobium meliloti*, and Fung-LCOs molecules commonly produced by a vast number of rhizospheric fungi (Rush *et al*., 2020). We carried out two Genome-Wide Association Studies (GWAS) within a collection of 173 accessions of *M. truncatula* (Bonhomme *et al*., 2014), whose seedlings have been either treated with cognate Nod-LCOs, mainly LCO-IV(C16:2, Ac, S) or with the Fung-LCOs, LCO-V(C18:1, Fuc/MeFuc) (Rush *et al*., 2020). By doing so, we could compare root responses to Nod- and Fung-LCOs in a way that is not possible using the reference A17 genotype and uncovered specific genetic determinants underlying these root responses. These results shed light on how legumes can cope with rhizospheric structurally related signals emitted by distinct microbes.

## Materials and Methods

### Production of lipo-chitooligosaccharide molecules

The Fung-LCOs used here were LCO-V(C18:1, Fuc/MeFuc) synthesized by metabolically engineered *Escherichia coli* as described in (Samain *et al*., 1997; Samain *et al*., 1999; Ohsten Rasmussen *et al*., 2004; Chambon *et al*., 2015), the fucosyl and methylfucosyl substitutions on the reducing end were obtained as described in (Djordjevic *et al*., 2014). They were chosen as they are the most representative of the fungal LCOs (Rush *et al*., 2020). *Sinorhizobium meliloti* Nod factors, named thereafter “Nod-LCOs” [mainly LCO-IV(C16:2, Ac, S)] were extracted from *S. meliloti* culture supernatants by butanol extraction, and purified by high-performance liquid chromatography (HPLC) on a semi-preparative C18 reverse phase column, as described in (Roche *et al*., 1991b). Nod-LCO and Fung-LCO structures (Fig. S1) were verified by mass spectrometry as described in (Cope *et al*., 2019).

### Plant material, experimental design and root phenotyping

A collection of 173 *M. truncatula* accessions (http://www.medicagohapmap.org) provided by the INRAE *Medicago truncatula* Stock Center (Montpellier, France; www1.montpellier.inra.fr/BRC-MTR/), was used for phenotyping experiments. These accessions are representative of the overall genetic diversity of *M. truncatula* and belong to the CC192 core collection (Ronfort *et al*., 2006). GWAS for various phenotypic traits have already been performed using this collection (Stanton-Geddes *et al*., 2013; Bonhomme *et al*., 2014; Yoder *et al*., 2014; Kang *et al*., 2015; Bonhomme *et al*., 2019).

*M. truncatula* seeds were scarified with sulfuric acid, sterilized in bleach (2.5%) for four minutes, washed in sterile water, and transferred on sterile agar plates for 2.5 days in the dark at 15°C to synchronize germination. Seedlings were then grown *in vitro* on square Petri dishes (12×12 cm) under 16 h light and 8 h dark at 22°C, with a 70° angle inclination, on modified M-medium as described in (Bonhomme *et al*., 2014). This medium contained either (i) the “Nod” treatment in which Nod-LCOs were incorporated at a concentration of 10^−8^ M, (ii) the “Fung” treatment in which Fung-LCOs, less water soluble than the sulfated Nod-LCOs, were incorporated at a concentration of 10^−7^ M to ensure a final experimental concentration close to 10^−8^ M (Ohsten Rasmussen *et al*., 2004), and (iii) two control (CTRL) conditions where acetonitrile 50% was diluted 1000x (CTRL-Fung) and 10000x (CTRL-Nod). Each accession of *M. truncatula* was phenotyped in two independent biological repeats, with 15 seedlings per repeat (5 seedlings per plate), for each treatment (Nod, Fung, CTRL-Nod, CTRL-Fung).

For each treatment, the lateral root number (LR) of each seedling was followed at four time points of plant development: 5, 8, 11 and 15 days after seedling transfer on LCO-containing medium. In addition, the primary root length (RL) was measured 5- and 11-days post treatment in order to calculate the lateral root density (LRD, *i*.*e*. the ratio of the lateral root number over the primary root length of each plant). All these measurements were carried out using the image analysis software Image J, using scans of plates. In order to summarize the kinetics of lateral root number appearance over the four time points, we calculated for each plant the Area Under the Lateral Root Progress Curve -AULRPC- (Fig. S2) using the R statistical package “agricolae”. Overall, nine phenotypic variables were recorded for each plant and for each treatment: LR_5d, LR_8d, LR_11d, LR_15d, RL_5d, RL_11d, LRD_5d, LRD_11d and AULRPC.

### Statistical modeling of phenotypic data

For the Nod and Fung treatments separately, as well as for the control of each treatment (i.e. mock treated plants of the Nod- or Fung-LCOs experiments), adjusted means of each accession (coefficients) were estimated for each of the nine phenotypic variables by fitting the following linear model with fixed effects: y_*ijk*_ = accession_*i*_ + repeat_*j*_ + ε_*ijk*_, where y_*ijk*_ is the phenotypic value of the *k*th plant of the *j*th repeat for the *i*th accession. Since variation in the root system development naturally occurred within and among accessions both in control and Nod/Fung-treated plants, for LR, RL, LRD and AULRPC variables, an additional variable of induction/repression of the root system development was estimated for each accession by subtracting the coefficient value under treatment with Nod- or Fung-LCOs by the coefficient value under control condition (i.e. CTRL-Nod or CTRL-Fung). GWAS was performed using these variables, referred to as “delta”, estimated for each accession on Nod and Fung-LCOs treatments separately (delta_LR_5d, delta_RL_5d, delta_LRD_5d, delta_LR_8d, delta_LR_11d, delta_RL_11d, delta_LRD_11d, delta_LR_15d, delta_AULRPC).

### Association mapping and local score analyses of phenotypic data

GWAS was performed on the phenotypic variables described in the previous section, based on phenotypic values for 173 accessions of *M. truncatula*. We used the Mt4.0 Medicago genome and SNP version to perform GWAS (see http://www.medicagohapmap.org/). A set of 5,165,380 genome-wide SNPs was selected with a minor allele frequency of 5% and at least 90% of the 173 accessions scored across the *M. truncatula* collection. The statistical model used for GWAS was the mixed linear model (MLM) approach implemented in the EMMA eXpedited (EMMAX) software (Kang *et al*., 2010). The MLM is used to estimate and then test for the significance of the allelic effect at each SNP, taking into account the genetic relationships between individuals to reduce the false positive rate. Genetic relationships among accessions were estimated using a kinship matrix of pairwise genetic similarities which was based on the genome-wide proportion of alleles shared between accessions, using the whole selected SNP dataset.

The MLM first implements a variance component procedure to estimate the genetic (σ^2^_a_) and residual (σ^2^_e_) variances from the variance of the phenotypic data, by using the kinship matrix in a restricted maximum likelihood framework. Narrow-sense heritabilities (*i*.*e*. portion of the total phenotypic variation attributable to additive genetic effect, h^2^_ss_) of each phenotypic variable were calculated from estimates of σ^2^_a_ and σ^2^_e_. For each marker a Generalized Least Square *F*-test is used to estimate the effects β_k_ and test the hypothesis β_k_ *=* 0 in the following model: *y*_i_ = β_0_ + β_k_X_ik_ + η_i_, with X_ik_ the allele present in individual *i* for the marker *k*, and η_i_ a combination of the random genetic and residual effects (Kang *et al*., 2010). As in previous GWAS in *M. truncatula* (Bonhomme *et al*., 2014; Rey *et al*., 2017), we used a genome-wide 5% significance threshold with Bonferroni correction for the number of blocks of SNPs in linkage disequilibrium (*i*.*e. p*-value ≤ 10^−6^), to identify significant associations following the *F*-test on the estimated allele effect size at each SNP.

In order to detect small-effect QTL that would not pass the 10^−6^ significance threshold, we performed a local score approach (Fariello *et al*., 2017; Bonhomme *et al*., 2019) on SNP *p*-values. The local score is a cumulative score that takes advantage of local linkage disequilibrium (LD) among SNPs. This score, defined as the maximum of the Lindley process over a SNP sequence (*i*.*e*. a chromosome), as well as its significance threshold were calculated based on EMMAX *p*-values, using a tuning parameter value of ξ = 3, as suggested by simulation results (Bonhomme *et al*., 2019). R scripts used to compute the local score and significance thresholds are available at https://forge-dga.jouy.inra.fr/projects/local-score/documents.

## Results

### Natural variation in the stimulation of lateral root formation by Fung- and Nod-LCOs in *M. truncatula*

The Fung-LCOs molecules used in this study belong to the class of LCOs most commonly found in fungi (Rush *et al*., 2020). They are LCO-V(C18:1, Fucosylated/MeFucosylated). On the other hand, the Nod-LCOs are specific to the rhizobium *S. meliloti* (Roche *et al*., 1991b) that nodulates *M. truncatula*. These Nod-LCOs are mainly LCO-IV(C16:2, Ac, S). The LCOs used therefore display some structural commonalities but also some specificities (see Fig. S1).

Growth of the 173 accessions of *M. truncatula* in the presence of Fung-LCOs or Nod-LCOs led to 67% and 87% of them with delta_AULRPC values above 0, respectively. This suggests a global trend of LCO stimulation of lateral root formation (LRF), especially with Nod-LCOs (Fig. 1a,b). This trend appeared early in the experiment since LRF was stimulated in 72% and 83% of the accessions 5 days following Fung-LCO and Nod-LCO treatments, respectively (Table 1). Among these accessions, the reference genotype A17 was strongly stimulated by Nod-LCOs over the time course, but not by Fung-LCOs (Fig. 1a,b). Since LRF stimulation showed substantial variation across the *M. truncatula* collection, we estimated the heritability, namely the proportion of phenotypic variation observed that was due to genetic variation in the collection (Table 1). In response to Fung-LCOs, the heritability was relatively low (h^2^_ss_ ≤ 0.16) for phenotypic variables quantifying variation in lateral root (LR) number and density, and showed a clear tendency to increase over time (h^2^_ss_ = 0.16 for LR number at 15 days post treatment and h^2^_ss_ = 0.15 for LR density at 11 days). In contrast, in response to Nod-LCOs the heritability of variation in lateral root number and density was strong at early times (i.e. 0.66 and 0.75 at 5 days post treatment, respectively) and decreased over time but remained relatively high (i.e. > 0.22 and 0.35, respectively). Interestingly, variation of primary root length in response to Fung- and Nod-LCOs was also observed. Its heritability was stronger for Nod-LCOs at 11 days (h^2^_ss_ = 0.36, Table 1). In the case of treatment with Nod-LCOs, these results indicate that variation in LRF stimulation, but also in primary root length stimulation, was largely due to genetic variation in the collection, especially at early steps, showing the importance of natural variation in the genetic control of LRF and primary root length stimulation by Nod-LCOs in *M. truncatula*. In the case of treatment with Fung-LCOs, however, the strong level of LRF stimulation as well as the low heritability at early steps (0 ≤ h^2^_ss_ ≤ 0.06, see Table 1) support the hypothesis that the root response to Fung-LCOs in *M. truncatula* is much more genetically constrained than the root response to Nod-LCOs.

**Table 1.**
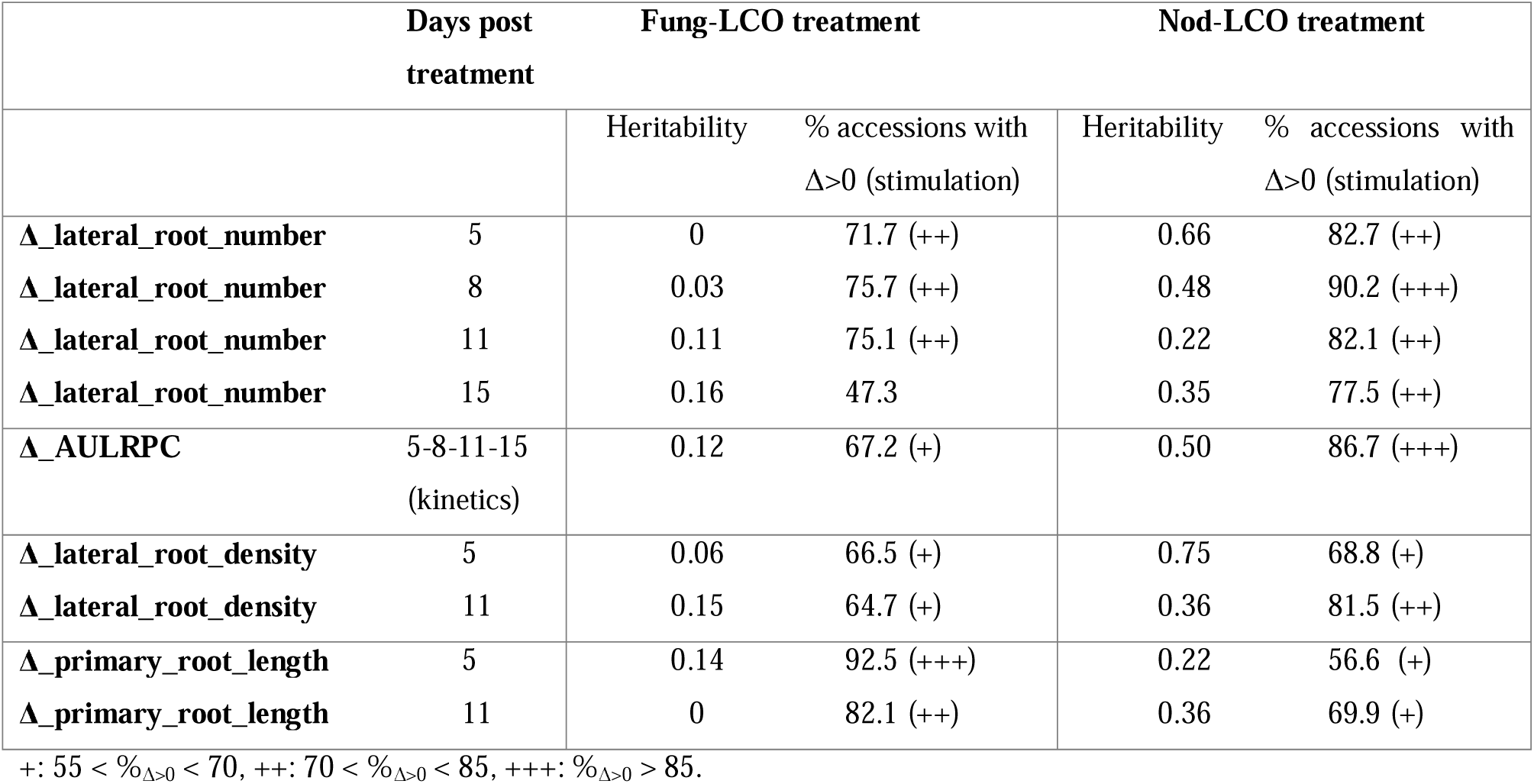
Estimation of narrow-sense heritability for different phenotypic variables measuring lateral root stimulation.

**Figure 1.**
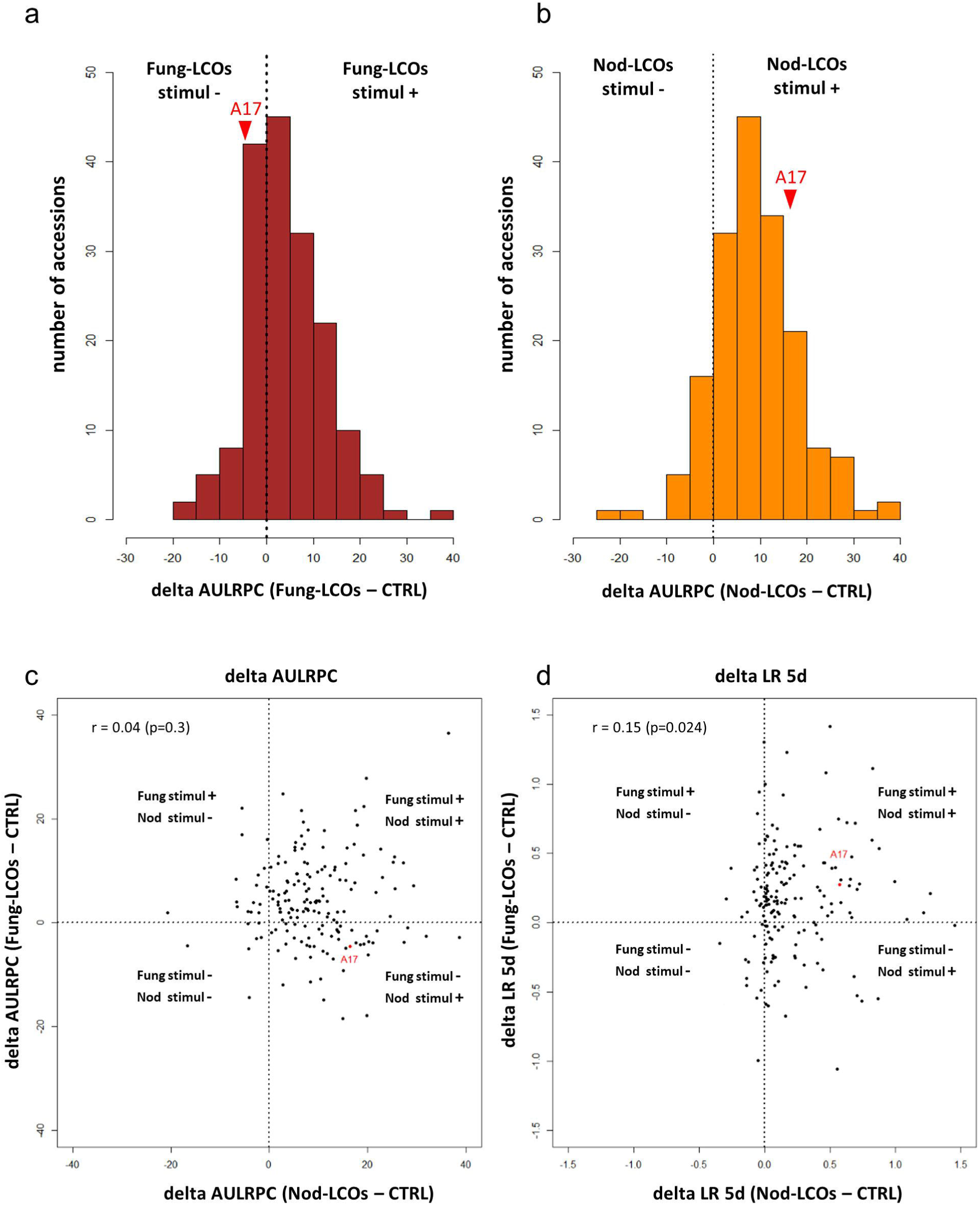
*Medicago truncatula* stimulation of root development by Fung- and Nod-LCOs. Quantitative variation in the stimulation of root development is observed in response to (a) Fung- and (b) Nod-LCOs, with 67% and 87% of the 173 accessions of *M. truncatula* showing stimulation of root development, respectively. This root development was measured for 15 days and expressed as the delta_AULRPC (see Fig. S2). The position of the reference genotype A17, relative to the other accessions, is indicated by a red arrow head. (c) Plot of delta_AULRPC (Nod-LCOs – CTRL) values versus delta_AULRPC (Fung-LCOs – CTRL) values and (d) plot of delta_LR_5d (Nod-LCOs – CTRL) versus delta_LR_5d (Fung-LCOs – CTRL) values for 173 accessions of *Medicago truncatula*, indicating a weak correlation between the stimulation by Fung- and Nod-LCOs. Vertical and horizontal dashed lines indicate equal states of root development between treatment (Fung- or Nod-LCOs) and control conditions (CTRL). The reference genotype A17 is indicated in red.

Since Fung and Nod-LCOs show a high structural homology and both stimulated LRF in most genotypes, we tested whether accessions highly stimulated by Nod-LCOs were also highly stimulated, not stimulated or even repressed by Fung-LCOs. Interestingly, for all variables, we found no correlation between the stimulations by Fung- and Nod-LCOs, except at 5 days where we found a significant but weak positive correlation for the variation in lateral root number (r = 0.15, *p*-value = 0.024). The lack of global correlation between LRF stimulation by Fung-LCOs and LRF stimulation by Nod-LCOs is illustrated in (Fig. 1c,d), with the delta_AULRPC variable which captures root development over time, and with the lateral root number at 5 days (delta_LR_5d) which captures early steps of root development.

Overall, these results suggest that (i) both Fung- and Nod-LCOs have the property to stimulate LRF in a quantitative manner, and (ii) genetic variation seems more influential in the root response to Nod-LCOs than to Fung-LCOs. To better understand the genetic determinants underlying these contrasted phenotypic responses, we performed a Genome-Wide Association Study.

### Genetic determinants underlying quantitative variation in root responsiveness to Fung- and Nod-LCOs in *M. truncatula*

GWAS was performed separately for Fung and Nod-LCO treatments, for each of the nine phenotypic variables measuring: (i) variation of the lateral root number (delta_LR_5d, delta_LR_8d, delta_LR_11d and delta_LR_15d), (ii) lateral root density (delta_LRD_5d and delta_LRD_11d), (iii) primary root length (delta_RL_5d, delta_RL_11d) and (iv) lateral root progress curve (delta_AULRPC) over time (5, 8, 11 and 15 days). Across all phenotypic variables measured in response to Fung-LCOs and Nod-LCOs, *p-value*-based tests performed using EMMAX respectively identified 24 and 70 genomic regions or loci significant at the *p*-value threshold of 10^−6^. Using the local score approach, more significant candidate genomic regions were identified as associated with root response to Fung- and Nod-LCOs, respectively 71 and 123 loci and 1 common locus (Table S1). All the loci identified with the EMMAX approach are nested within the local score results. Identified loci contain 1 to 11 genes, corresponding to 291 possible genes in total (see Table S1).

A global view of the genome-wide quantitative genetic bases of LRF stimulation kinetics following treatment with LCOs could be obtained by the local score analysis of the delta_AULRPC variable (Fig. 2a, b). Genetic variation involved in LRF stimulation specifically in response to Fung-LCOs mainly relied on four candidate loci; a gibberellin 2-oxidase (*Medtr1g086550*, GA2OX) and three receptor-like kinases: a putative Feronia receptor-like kinase - *Medtr6g015805*-, a crinkly 4 receptor like kinase CCR4-like protein - *Medtr3g464080* -, and a Serine/Threonine kinase PBS1 - *Medtr8g063300* – (Fig. 2a, Table S1). One major locus on chromosome 7, containing genes from the leguminosin LEED.PEED family (Trujillo *et al*., 2014), but also kinase encoding genes with potential carbohydrate-binding properties were specifically involved in response to Nod-LCOs (Fig. 2b, Table S1). Only one candidate genomic region involved in the response to Fung-LCOs and Nod-LCOs was identified in this study, by the GWAS analysis of delta_AULRPC and primary root length (delta_RL_5d) phenotypic variables (Table S1). This region on chromosome 8 contains three genes among which two encode “embryonic abundant protein”, annotated as BURP domain-containing protein by the new *M. truncatula* genome version Mt5 (Pecrix *et al*., 2018).

**Figure 2.**
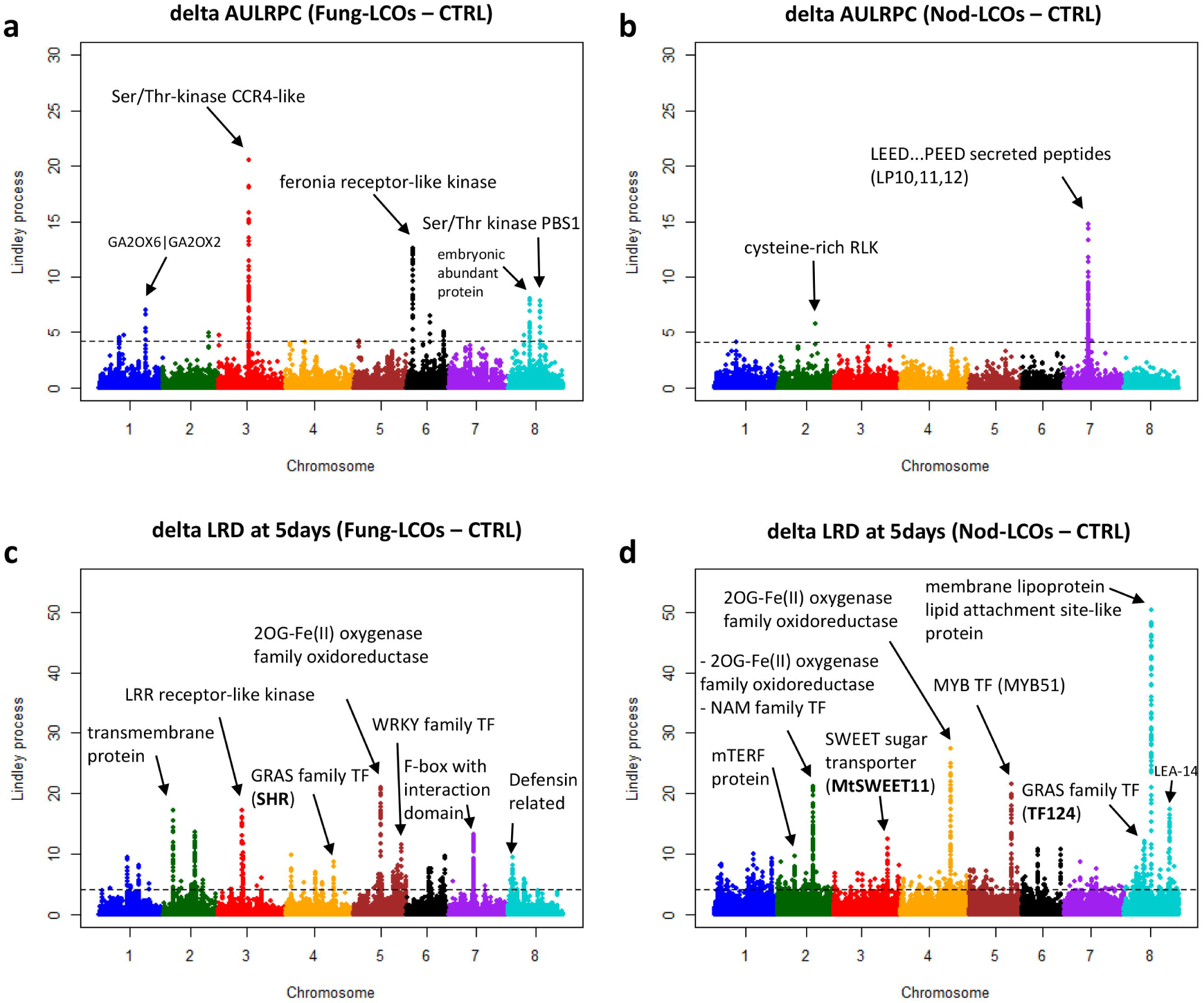
GWAS results using a local score approach on *Medicago truncatula* stimulation of lateral root development by Fung- and Nod-LCOs. Each Manhattan plot shows on the y-axis the Lindley process (the local score with the tuning parameter ξ = 3) for SNPs along the eight chromosomes (x-axis), with the dashed line indicating the maximum of the eight chromosome-wide significance thresholds. The local score is shown for GWAS of four phenotypic variables: (a) delta_AULRPC (Fung-LCOs – CTRL), (b) delta_AULRPC (Nod-LCOs – CTRL), (c) delta_LRD_5d (Fung-LCOs – CTRL) and (d) delta_LRD_5d (Nod-LCOs – CTRL). The most significant candidate genes and their predicted functions are indicated by arrows on the plots (see Table S1).

A more precise view of the genome-wide quantitative genetic bases of the early steps of LRF stimulation following treatment with LCOs could be obtained by the local score analysis of the delta_LRD_5d variable (Fig. 2c, d). Interestingly, this phenotypic variable showed highly contrasted heritability values between treatments with Fung- and Nod-LCOs (h^2^_ss_ = 0.06 and 0.75, respectively; Table 1). Among 34 candidate genomic regions identified in response to Fung-LCOs, we identified four highly significant candidate genes whose predicted proteins show good homology for known functions, such as a dioxygenase (*Medtr5g055800*), an LRR receptor-like kinase (*Medtr3g452970*), a WRKY family transcription factor (*Medtr5g091390*) and a GRAS family transcription factor (*Medtr4g097080*) whose homolog in *Arabidopsis thaliana* is SHORT-ROOT -SHR- (Helariutta *et al*., 2000). Among 49 candidate genomic regions identified in response to Nod-LCOs for the delta_LRD_5d variable, we identified 4 highly significant candidate genes, among which two encoded dioxygenases (*Medtr4g100590, Medtr2g068940*), one MYB transcription factor (*Medtr5g081860*, MYB51) and the most significant one encoding a putative membrane lipoprotein lipid attachment site-like protein (*Medtr8g464760*), annotated as thioredoxin-like protein in Mt5 genome. This analysis also detected two known genes encoding a sugar transporter (*Medtr3g098930*, MtSWEET11) and a GRAS family transcription factor (*Medtr8g442410*, TF124) (Fig. 2d).

### Combination of GWAS results with Gene ontology classification highlights enrichment in signaling functions

GWAS most significant genes can give a first hint to determine some of the mechanisms involved in root response to LCOs. However local score also highlighted minor QTL/genes and allowed us to identify several dozen of supplementary genes. To gain further insights from these data, we performed a Gene Ontology (GO) enrichment analysis using the Medicago Superviewer interface (Herrbach *et al*., 2017) (Fig. 3a,b). 71 and 134genes identified in the Fung-LCO and Nod-LCO GWAS were classified, respectively. At the “biological process” level, both the Nod and Fung-LCO datasets were enriched in biological functions related to “other metabolic processes”. The Nod-LCO data were also enriched in transcription related biological processes. Although the Fung-LCO data did not show any significant enrichment in transcription function at the “biological process” level, they were, as the Nod-LCO data, enriched in transcription factor and kinase activities at the “molecular function” level (Fig. 3 a,b). This is in accordance with the numerous loci associated with Receptor-like kinases or transcription factors (TF) found in both datasets (see Table S1). Accordingly, the Nod-LCO data showed enrichment in nuclear and plasma membrane associated “cellular component” (Fig. 3b). Many of the metabolic functions from the Nod-LCO candidates and of the genes underlying the “protein metabolism” biological process enriched with Fung-LCOs were associated with phosphorylation, so possibly also with signaling pathways. In addition, a significant proportion of loci were associated with oxido-reduction processes and cell-wall metabolism enzymes (pectin-esterases, cellulose synthase, phenylalanine ammonia-lyase-like protein). Although not specifically enriched in these datasets, we also found several hormone related genes. For instance, auxin signaling (AUX/IAA and Auxin Response Factor, ARF) and auxin transport (efflux carriers) genes were found in the Nod-LCO data whereas an ethylene receptor and an ethylene responsive TF were found in the Nod-LCO and Fung-LCO data, respectively (Table S1).

**Figure 3.**
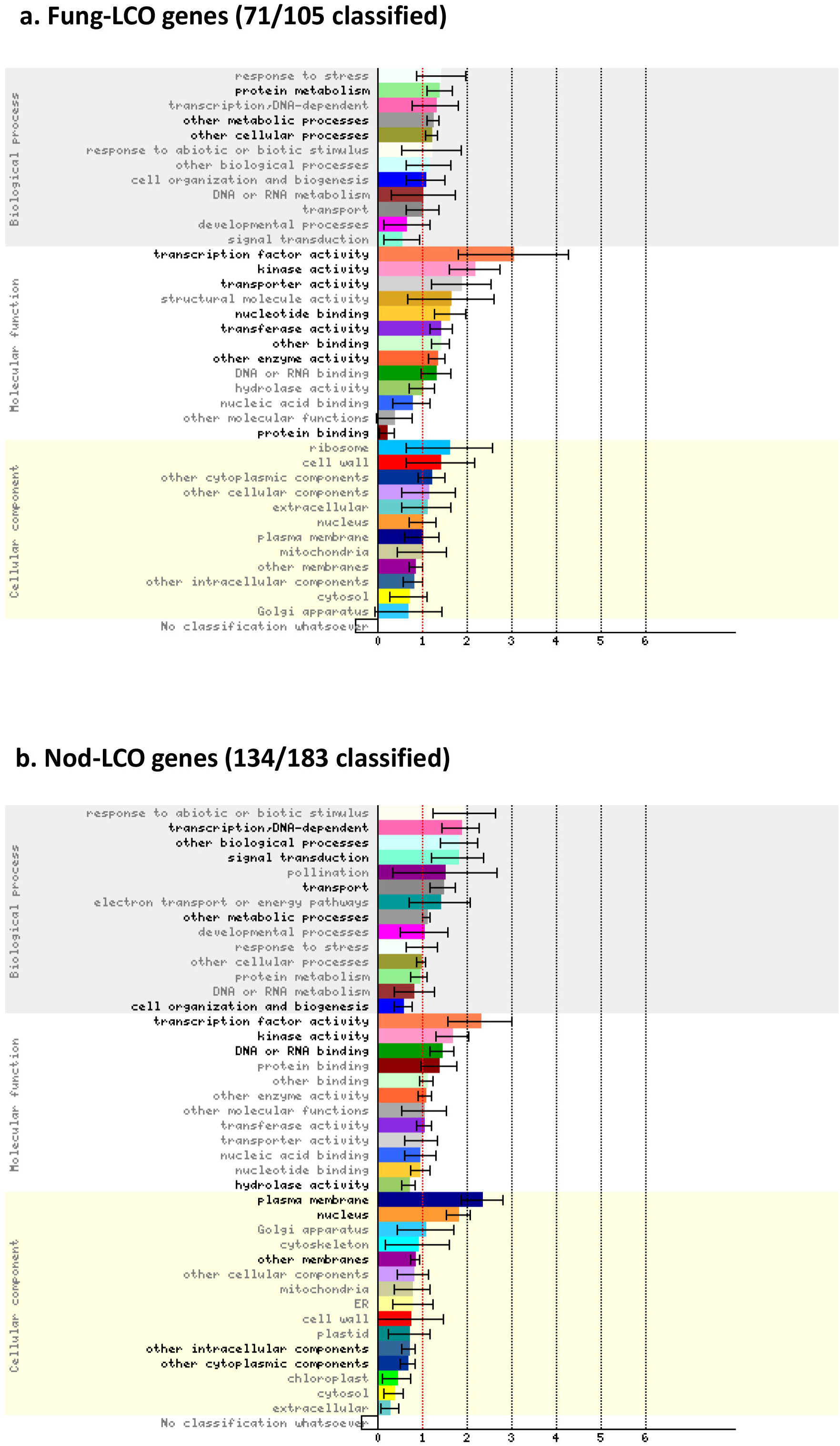
Gene ontology enrichment for the Nod-LCOs and Fung-LCOs candidate loci identified by GWAS (local score results) in *Medicago truncatula*. Graphical summary of the gene ontology (GO) classification ranking of Fung-LCO candidate genes (a, 71/105 represented) and Nod-LCO candidate genes (b, 134/183 represented) using the Classification SuperViewer tool from bar.utoronto.ca adapted to *Medicago truncatula*. Bars represent the normed frequency of each GO category for the given sets of genes compared to the overall frequency calculated for the Mt4.0 *Medicago truncatula* (see Herrbach *et al*., 2017). Hence, a ratio above 1 means enrichment and below 1 means under-representation. Error bars are standard deviation of the normed frequency calculated by creating 100 gene sets from the input set by random sampling and computing the frequency of classification for all of those data sets across all categories. Hypergeometric enrichment tests on the frequencies were performed and GO categories showing significant *p*-values (< 0.05) are printed bold. GO categories are displayed for each GO subclass ranked by normed frequency values.

To gain further insight in possible biological processes where those loci could be involved, we also compiled transcriptional expression data from the literature and the knowledge database LEGOO (Carrère *et al*., 2020). Data could be obtained for 148 out of the 291 candidate genes and are summarized in Table S2. As expected, a majority of genes were found in symbiotic studies (nodulation or mycorrhization, 123 genes) or with LCO treatments (25 genes among which 23 are also found in the symbiotic data). However, available expression data was not restricted to these symbiotic interactions. Indeed, expression data could also be retrieved from nitrate or phosphate starvation experiments or from data obtained with Medicago root pathogens or defense elicitors (Table S2).

## Discussion

In this study, we asked whether a legume, here *M. truncatula*, is capable of distinguishing lipo-chitooligosaccharide molecules that share similar structures and induce the same developmental root responses. Regulation of root development by LCOs seems to be a conserved plant response observed in legume and non-legume plants (Sun *et al*., 2015; Tanaka *et al*., 2015; Buendia *et al*., 2019), raising the question of its possible evolutionary origin and molecular conservation. The Nod-LCO molecules we used, LCO-IV(C16:2, Ac, S), are produced by the rhizobial symbiont of *M. truncatula*. These LCOs can be considered as very specific symbiotic signals, with a key role in the narrow host specificity that characterizes the rhizobium legume symbiosis (RLS). The simple absence of the sulfate group on the reducing end of the Nod-LCOs renders them inactive symbiotically on Medicago (Roche *et al*., 1991a; Bensmihen *et al*., 2011). In contrast, the Fung-LCO molecules used here, LCO-V(C18:1, Fuc/MeFuc), are not only a form of LCOs commonly found in AM fungi, but they can also be produced by pathogenic or saphrophytic fungi (Rush *et al*., 2020) and can thus be considered as a common, almost universal, hallmark of fungal presence. Furthermore, it is worth noticing that even *Bradyrhizobia* and *Sinorhizobium* symbionts of soybean also produce LCO-V(C18:1, Fuc/MeFuc) (D’Haeze & Holsters, 2002; Wang *et al*., 2018), making them also non cognate Nod-LCO signals. By studying the ability of *M. truncatula* plants to respond to specific (Nod-LCOs) or wide-spread (Fung-LCOs) LCOs, we were thus considering a common situation encountered by plants in their natural environment where they must distinguish different LCO-producing microorganisms.

Here, we have exploited the large genetic diversity among *M. truncatula* natural accessions using a GWAS approach to compare the genetic bases underlying root developmental responses. The root phenotypic traits that we used, lateral root formation and lateral root density, were chosen because in the *M. truncatula* A17 reference accession these traits are stimulated by Nod factors and by the Myc-LCOs originally detected in AM fungi (Fig. S1) (Olah *et al*., 2005; Maillet *et al*., 2011). To address LR density, we also looked at primary root growth, a parameter that was not previously described as affected by Nod-LCOs in A17. Moreover, these traits are relatively easy to score, which was convenient to phenotype many accessions of *M. truncatula*.

### The Fung-LCO structures stimulate root development of *M. truncatula* in a quantitative way

Our results clearly show that the Fung-LCO molecules tested, LCO-V (C18:1, Fuc/MeFuc) can also stimulate LRF in *M. truncatula*. This LRF stimulation is variable among the accessions, and the trait would have been missed if we had only studied the reference accession, A17, which is poorly responsive (Fig. 1), as previously shown with *Sinorhizobium fredii* Nod factors, LCO-V (C18:1, MeFuc) (Olah *et al*., 2005). Also, in contrast to what was previously observed in A17 (Olah *et al*., 2005), we could detect some positive effect of Nod-LCOs on primary root length, especially at later time points (11 days). The majority of accessions responded positively to Fung-LCOs for this growth parameter at both 5 and 11 days. Accordingly, we found a number of loci associated with the variation in primary root length phenotype (Table S1). This underlines the power of the natural variation approach that can detect more responsive genetic backgrounds and reveal new genetic determinants that would have passed unnoticed in forward and reverse genetic screens with classical reference accessions. Similarly, GWAS results obtained on root architecture modification of *Arabidopsis thaliana* upon hormonal treatments identified that the Col-0 reference accession is not the most responsive to auxin (Ristova *et al*., 2018).

### *Medicago truncatula* can distinguish between Fung-LCOs and Nod-LCOs

The lack of overlap, with only one exception and for different parameters, between the loci identified in the Nod-LCO and Fung-LCO GWAS is striking. This lack of overlap is consistent with the weak correlation between the ability of one accession to respond to Nod- and to Fung-LCOs (Fig. 1). The absence of common genes (except one locus) highlighted in the two GWAS, and the very different heritability values found associated with the Fung-LCO and Nod-LCO responses, indicate that *M. truncatula* clearly distinguishes these signals, although they have similar structures and cause the same root response. This can be due to specific receptors (no data is available yet concerning plant receptors for the Fung-LCOs we used) and/or to divergence in downstream signaling pathways. The latter hypothesis is consistent with the enrichment in signaling functions we observed in the GWAS genes (Fig. 3). Nod-LCO and Myc-LCO stimulation of LRF requires the CSSP in *M. truncatula* (Olah *et al*., 2005; Maillet *et al*., 2011). However, previous transcriptomic studies performed with Myc-LCO structures which are closer to those of Nod-LCOs from *S. meliloti* (Fig. S1) identified that Myc-LCO signaling can also act independently of the CSSP gene *MtDMI3* (Czaja *et al*., 2012; Camps *et al*., 2015). It would be interesting to test whether the Fung-LCOs we used here require signaling from the CSSP to activate the LRF responses in *M. truncatula*. CSSP mutants are available in the *M. truncatula* A17 genetic background but this accession is poorly responsive to these new Fung-LCOs in our assays (see Fig. 1).

### Genetic determinants of *M. truncatula* responses to Fung-LCOs and Nod-LCOs

#### Cell wall, root growth and developmental signaling pathways associated loci

Only one of the genes or loci identified in the two GWAS analyses was found to be common. This region contained two genes annotated as BURP domain-containing proteins, which define a group of proteins specific to plants (Table S1). This domain was named from the four members of the group initially identified, BNM2, USP, RD22, and PG1beta and is commonly found in plant cell wall proteins (Hattori *et al*., 1998; Wang *et al*., 2015). Cell-wall related functions, like-cell-wall remodeling, could be linked to root growth promotion activities of the LCO molecules, and additionally might be related to the root hair deformation capacities of LCOs (Esseling *et al*., 2003). One gene associated with this locus (*Medtr8g046000*) was previously described as down-regulated by Nod-LCOs in the root epidermis (4h after 10^−8^M Nod-LCO treatment) (Jardinaud *et al*., 2016), downregulated in nodules at 4 and 10 dpi, compared to roots (El Yahyaoui *et al*., 2004) and upregulated in roots mycorrhized with *Rhizophagus irregularis* at 28 dpi compared to non-mycorrhizal control roots (Hogekamp *et al*., 2011) (see Table S2).

In the Fung-LCO GWAS, we found some signaling genes that could have a role in LRF. These are the receptor like kinase (RLK) CRINKLY 4 (CCR4) (*Medtr3g464080*), and a GRAS TF (*Medtr4g097080*) related to the *SHORTROOT* gene of *Arabidopsis*, known to control root development (Helariutta *et al*., 2000; De Smet *et al*., 2008), although neither of these two genes has been characterized in *M. truncatula*. Among the putative RLK genes detected in the Fung-LCO GWAS, there was also one that could encode a Feronia RLK (*Medtr6g015805)*. Interestingly, this protein regulates root growth of *A. thaliana* (Haruta *et al*., 2014) but also plant immune signaling by sensing cell-wall integrity (Stegmann *et al*., 2017), two biological processes also regulated by LCOs. Similarly, we identified several receptor-like cytosolic kinases (RLCKs), also known as PBS1-like kinases, from the subfamily VII in the Nod-LCO data. Some genes from this subfamily are involved in PAMP-triggered immunity (PTI), including chitin responses in *A. thaliana* (Rao *et al*., 2018).

#### Phytohormone associated loci

Relatively few hormone-related genes were identified in the two GWAS and they were all different. The ethylene-related genes *Medtr1g069985* and *Medtr1g073840* were found in Fung-LCO and Nod-LCO GWAS, respectively. A gibberellin-related GA2 oxidase gene (*Medtr1g086550*) and a few auxin transporter genes (*Medtr5g024530, Medtr5g024560* and *Medtr5g024580*) were found in the Fung-LCO and Nod-LCO GWAS, respectively. GA2 oxidase is predicted to be a catabolic enzyme that degrades gibberellins (GA) (Yamaguchi, 2008). In *M. truncatula*, in contrast to Arabidopsis, GAs are negative regulators of LRF (Fonouni-Farde *et al*., 2019). They are also negative regulators of nodulation and mycorrhization (Foo *et al*., 2013; Bensmihen, 2015) so down regulation of the GA content could stimulate LRF, nodulation and mycorrhization. Interestingly, all the auxin-related functions were found in the Nod-LCO GWAS only. This could be related to the tight developmental links between LR formation and nodule organogenesis and their common need for auxin accumulation in *M. truncatula* (Schiessl *et al*., 2019; Soyano *et al*., 2019).

#### Endosymbiosis associated loci

Several other loci we identified could also be related to symbiosis. When comparing with previous transcriptomic studies, we found 123 genes (78 for Nod-LCOs, 44 for Fung-LCOs and one found in both studies) expressed during symbiotic processes (nodulation or mycorrhization, Table S2). This represents an important overlap probably linked to the role of these molecules as pre-symbiotic or symbiotic signals to prepare for specific symbiotic events. We could even find some very specific LEED…PEED loci that are only expressed in nodules (Trujillo *et al*., 2014). Along the same line, *MtSWEET11* (found for the difference in LRD at 5 days with Nod-LCOs, Table S1) was previously shown to be expressed in infected root hairs, and more specifically in infection threads and symbiosomes during nodulation in *M. truncatula*. However, knock out of this gene did not impair RLS, possibly due to genetic redundancy (Kryvoruchko *et al*., 2016). This illustrates the interest of GWAS to identify genes without any redundancy issues. Some genes identified in our Nod-LCO GWAS were also found in a previous GWAS of nodulation. For example, *Medtr1g064090/Medtr1te064120* (annotated as a phenylalanine ammonia-lyase-like protein / Copia-like polyprotein/retrotransposon) and *Medtr2g019990* (annotated as a Serine/Threonine-kinase PBS1-like protein) were previously found by Stanton-Geddes and colleagues as associated with nodule numbers in the lower part of the root (Stanton-Geddes *et al*., 2013). Two other loci *Medtr3g034160* (galactose oxidase) and *Medtr5g085100* (AP2 domain class transcription factor) were respectively found as associated with nodule numbers in the upper part of the root and with strain occupancy in the lower part of the root (Stanton-Geddes *et al*., 2013).

We did not find any known CSSP or LysM-RLK genes among our loci detected by GWAS. This is somehow expected as constrained natural variability on these essential symbiotic genes due to selective processes was often found in previous nucleotide polymorphism analyses (De Mita *et al*., 2006; De Mita *et al*., 2007; Grillo *et al*., 2016) and in previous GWAS studies performed on nodulation phenotypes (Stanton-Geddes *et al*., 2013). This also suggests that these genes are not major determinants of natural variability in root developmental responses to LCOs, although some LysM-RLK genetic variants likely account for rhizobia host-specificity (Sulima *et al*., 2017; Sulima *et al*., 2019).

### Evolutionary origin of *M. truncatula* responses to Fung-LCOs and Nod-LCOs

Our GWAS results also raise interesting questions on the evolutionary origin of the root growth stimulation role of LCOs. Indeed, the two different LCO structures (from different microbial origins) triggered LRF stimulation on a high number of Medicago accessions. The low heritability of plant responses to Fung-LCOs (with a maximum of 0.16 for the difference in LR number at 15 days), compared to that of plant responses to Nod-LCOs (with a maximum of 0.75 for lateral root density at 5 days) is not due to a lack of activity of the Fung-LCOs since 67% to 76% of the accessions did show a positive root growth response to these LCOs. This rather suggests that the genetic determinants of the Fung-LCO responses are more “fixed” (*i*.*e*. less variable) than those of the Nod-LCO responses. The low genetic variability of responses to these widespread Fung-LCO structures is likely linked to their very ancient apparition in the fungi kingdom (Rush *et al*., 2020), and suggests that the ancient function(s) of these LCOs were non symbiotic. Ancient LCO functions could be LRF stimulation or the regulation of immunity in plants (Liang *et al*., 2013; Limpens *et al*., 2015; Feng *et al*., 2019), a function that may have predated the mycorrhizal symbiosis and has not been lost in Arabidopsis (Liang *et al*., 2014). LCOs could also be involved in other aspects of plant biology, yet to be discovered.

## Conclusion

In addition to providing many new genes potentially involved in regulating root development for future reverse genetic or allelic variant investigations, this study brings new evidence that plants can distinguish between specific and non-specific LCO signals and suggests that their recognition has had distinct evolutionary histories.

## Acknowledgments

This work was part of a program funded by the French Agence Nationale de la Recherche (ANR-14-CE18-0008, “NICE CROPS”). The authors thank the bioinformatics platform Toulouse Midi-Pyrenees (Genotoul). Thanks to V. Regard for help with formatting of TableS2. Mass spectrometry analyses were done with the support from the ICT-Mass Spectrometry and MetaToul-AgromiX facilities and from the MetaboHUB-ANR-11-INBS-0010 network. S.F. and S.C. received technical support of ICMG (FR 2607) mass spectrometry platform and partial financial support from the LABEX ARCANE and CBH-EUR-GS (ANR-17-EURE-0003), Glyco@Alps (ANR-15-IDEX-02), and PolyNat Carnot Institut (ANR-16-CARN-0025-01). This work was performed in the LRSV and LIPM (Toulouse, France), parts of the “Laboratoire d’Excellence” (LABEX) entitled TULIP (ANR-10-LABX-41).

## Author contributions

MB, SB: analyzed the data; MB, SB, CG, GB, CJ: wrote the manuscript; MB, GB, CG, CJ: designed the experiments; OA, EA, MG, FM, VPP: performed the experiments; SF, SC: synthesized the Fung-LCO molecules.

## Supplemental Figure legends

**Figure S1. Structures of the LCOs used in this study compared to the “original” Myc-LCOs as described in Maillet *et al*., 2011.**

The Fung-LCO molecules used in this study belong to the class of LCOs most commonly found in fungi (Rush *et al*., 2020): LCO-V(C18:1, Fucosylated/MeFucosylated). The Nod-LCOs used are specific to *S. meliloti*, rhizobial partner of *M. truncatula* (Roche *et al*., 1991b), mainly comprising LCO-IV(C16:2, Ac, S). As lipo-chitooligosaccharides, Fung-LCOs and Nod-LCOs have the same canonical structure but also differences such as their number of chitin residues (5 for Fung-LCOs and 4 for Nod-LCOs), their acyl chain on the non-reducing end (C18:1 for Fung-LCOs and C16:2 for Nod-LCOs) and their substituents on the reducing end (fucosyl or methylfucosyl for Fung-LCOs and sulfate for Nod-LCOs). The structures of the original Myc-LCOs described by Maillet et al.: LCO-IV(C16:0, S or C18:1, S) or LCO-IV(C16:0 or C18:1) (Maillet *et al*., 2011) are also shown for comparison.

**Figure S2. Lateral root formation phenotypic variables used in this study.**

Stimulation of *Medicago truncatula* root development with Fung- or Nod LCOs was monitored at different time-points (5, 8, 11 and 15 days post treatment), by counting lateral root number (LR), measuring primary root length (RL), calculating lateral root density (LRD, the ratio LR/RL) and measuring the Area Under the Lateral Root Progress Curve – AULRPC.

## References

Bensmihen S. 2015. Hormonal Control of Lateral Root and Nodule Development in Legumes. Plants (Basel) 4(3): 523–547.

Bensmihen S, de Billy F, Gough C. 2011. Contribution of NFP LysM Domains to the Recognition of Nod Factors during the Medicago truncatula/Sinorhizobium meliloti Symbiosis. PLoS One 6(11): e26114.

Bonhomme M, André O, Badis Y, Ronfort J, Burgarella C, Chantret N, Prosperi JM, Briskine R, Mudge J, Debéllé F, et al. 2014. High-density genome-wide association mapping implicates an F-box encoding gene in Medicago truncatula resistance to Aphanomyces euteiches. New Phytol 201(4): 1328–1342.

Bonhomme M, Fariello MI, Navier H, Hajri A, Badis Y, Miteul H, Samac DA, Dumas B, Baranger A, Jacquet C, et al. 2019. A local score approach improves GWAS resolution and detects minor QTL: application to Medicago truncatula quantitative disease resistance to multiple Aphanomyces euteiches isolates. Heredity (Edinb) 123(4): 517–531.

Buendia L, Maillet F, O’Connor D, van de-Kerkhove Q, Danoun S, Gough C, Lefebvre B, Bensmihen S. 2019. Lipo-chitooligosaccharides promote lateral root formation and modify auxin homeostasis in Brachypodium distachyon. New Phytol 221(4): 2190–2202.

Camps C, Jardinaud MF, Rengel D, Carrère S, Hervé C, Debellé F, Gamas P, Bensmihen S, Gough C. 2015. Combined genetic and transcriptomic analysis reveals three major signalling pathways activated by Myc-LCOs in Medicago truncatula. New Phytol 208(1): 224–240.

Carrère SB, Verdenaud M, Gough C, Gouzy JRM, Gamas P. 2020. LeGOO: An Expertized Knowledge Database for the Model Legume Medicago truncatula. Plant Cell Physiol 61(1): 203–211.

Catoira R, Galera C, de Billy F, Penmetsa RV, Journet EP, Maillet F, Rosenberg C, Cook D, Gough C, Dénarié J. 2000. Four genes of Medicago truncatula controlling components of a nod factor transduction pathway. Plant Cell 12(9): 1647–1666.

Chambon R, Despras G, Brossay A, Vauzeilles B, Urban D, Beau JM, Armand S, Cottaz S, Fort S. 2015. Efficient chemoenzymatic synthesis of lipo-chitin oligosaccharides as plant growth promoters. Green Chem. 17: 3923–3930.

Cope KR, Bascaules A, Irving TB, Venkateshwaran M, Maeda J, Garcia K, Rush TA, Ma C, Labbé J, Jawdy S, et al. 2019. The Ectomycorrhizal Fungus Laccaria bicolor Produces Lipochitooligosaccharides and Uses the Common Symbiosis Pathway to Colonize Populus Roots. Plant Cell 31(10): 2386–2410.

Czaja LF, Hogekamp C, Lamm P, Maillet F, Martinez EA, Samain E, Dénarié J, Küster H, Hohnjec N. 2012. Transcriptional responses toward diffusible signals from symbiotic microbes reveal MtNFP- and MtDMI3-dependent reprogramming of host gene expression by arbuscular mycorrhizal fungal lipochitooligosaccharides. Plant Physiol 159(4): 1671–1685.

D’Haeze W, Holsters M. 2002. Nod factor structures, responses, and perception during initiation of nodule development. Glycobiology 12(6): 79R–105R.

De Mita S, Santoni S, Hochu I, Ronfort J, Bataillon T. 2006. Molecular evolution and positive selection of the symbiotic gene NORK in Medicago truncatula. J Mol Evol 62(2): 234–244.

De Mita S, Santoni S, Ronfort J, Bataillon T. 2007. Adaptive evolution of the symbiotic gene NORK is not correlated with shifts of rhizobial specificity in the genus Medicago. Bmc Evolutionary Biology 7: 210.

De Smet I, Vassileva V, De Rybel B, Levesque MP, Grunewald W, Van Damme D, Van Noorden G, Naudts M, Van Isterdael G, De Clercq R, et al. 2008. Receptor-like kinase ACR4 restricts formative cell divisions in the Arabidopsis root. Science 322(5901): 594–597.

Djordjevic MA, Bezos A, Susanti, Marmuse L, Driguez H, Samain E, Vauzeilles B, Beau JM, Kordbacheh F, Rolfe BG, et al. 2014. Lipo-chitin oligosaccharides, plant symbiosis signalling molecules that modulate mammalian angiogenesis in vitro. PLoS One 9(12): e112635.

Dénarié J, Debellé F, Promé JC. 1996. Rhizobium lipo-chitooligosaccharide nodulation factors: signaling molecules mediating recognition and morphogenesis. Annu Rev Biochem 65: 503–535.

El Yahyaoui F, Kuster H, Ben Amor B, Hohnjec N, Puhler A, Becker A, Gouzy J, Vernie T, Gough C, Niebel A, et al. 2004. Expression profiling in Medicago truncatula identifies more than 750 genes differentially expressed during nodulation, including many potential regulators of the symbiotic program. Plant Physiol 136(2): 3159–3176.

Esseling JJ, Lhuissier FG, Emons AM. 2003. Nod factor-induced root hair curling: continuous polar growth towards the point of nod factor application. Plant Physiol 132(4): 1982–1988.

Fariello MI, Boitard S, Mercier S, Robelin D, Faraut T, Arnould C, Recoquillay J, Bouchez O, Salin G, Dehais P, et al. 2017. Accounting for linkage disequilibrium in genome scans for selection without individual genotypes: The local score approach. Mol Ecol 26(14): 3700–3714.

Feng F, Sun J, Radhakrishnan GV, Lee T, Bozsóki Z, Fort S, Gavrin A, Gysel K, Thygesen MB, Andersen KR, et al. 2019. A combination of chitooligosaccharide and lipochitooligosaccharide recognition promotes arbuscular mycorrhizal associations in Medicago truncatula. Nat Commun 10(1): 5047.

Fonouni-Farde C, Miassod A, Laffont C, Morin H, Bendahmane A, Diet A, Frugier F. 2019. Gibberellins negatively regulate the development of Medicago truncatula root system. Sci Rep 9(1): 2335.

Foo E, Ross JJ, Jones WT, Reid JB. 2013. Plant hormones in arbuscular mycorrhizal symbioses: an emerging role for gibberellins. Ann Bot 111(5): 769–779.

Genre A, Chabaud M, Balzergue C, Puech-Pagès V, Novero M, Rey T, Fournier J, Rochange S, Bécard G, Bonfante P, et al. 2013. Short-chain chitin oligomers from arbuscular mycorrhizal fungi trigger nuclear Ca2+ spiking in Medicago truncatula roots and their production is enhanced by strigolactone. New Phytol 198(1): 190–202.

Girardin A, Wang T, Ding Y, Keller J, Buendia L, Gaston M, Ribeyre C, Gasciolli V, Auriac M-C, Vernie T, et al. 2019. LCO Receptors Involved in Arbuscular Mycorrhiza Are Functional for Rhizobia Perception in Legumes. Current Biology 29(24):4249-4259.e5.

Gough C, Cullimore J. 2011. Lipo-chitooligosaccharide signaling in endosymbiotic plant-microbe interactions. Mol Plant Microbe Interact 24(8): 867–878.

Grillo MA, De Mita S, Burke PV, Solórzano-Lowell KL, Heath KD. 2016. Intrapopulation genomics in a model mutualist: Population structure and candidate symbiosis genes under selection in Medicago truncatula. Evolution 70(12): 2704–2717.

Haruta M, Sabat G, Stecker K, Minkoff BB, Sussman MR. 2014. A peptide hormone and its receptor protein kinase regulate plant cell expansion. Science 343(6169): 408–411.

Hattori J, Boutilier KA, van Lookeren Campagne MM, Miki BL. 1998. A conserved BURP domain defines a novel group of plant proteins with unusual primary structures. Mol Gen Genet 259(4): 424–428.

Helariutta Y, Fukaki H, Wysocka-Diller J, Nakajima K, Jung J, Sena G, Hauser MT, Benfey PN. 2000. The SHORT-ROOT gene controls radial patterning of the Arabidopsis root through radial signaling. Cell 101(5): 555–567.

Herrbach V, Chirinos X, Rengel D, Agbevenou K, Vincent R, Pateyron S, Huguet S, Balzergue S, Pasha A, Provart N, et al. 2017. Nod factors potentiate auxin signaling for transcriptional regulation and lateral root formation in Medicago truncatula. J Exp Bot 68(3): 569–583.

Hogekamp C, Arndt D, Pereira PA, Becker JD, Hohnjec N, Küster H. 2011. Laser microdissection unravels cell-type-specific transcription in arbuscular mycorrhizal roots, including CAAT-box transcription factor gene expression correlating with fungal contact and spread. Plant Physiol 157(4): 2023–2043.

Jardinaud MF, Boivin S, Rodde N, Catrice O, Kisiala A, Lepage A, Moreau S, Roux B, Cottret L, Sallet E, et al. 2016. A laser dissection-RNAseq analysis highlights the activation of cytokinin pathways by Nod factors in the Medicago truncatula root epidermis. Plant Physiol. 171(3):2256–76.

Kang HM, Sul JH, Service SK, Zaitlen NA, Kong SY, Freimer NB, Sabatti C, Eskin E. 2010. Variance component model to account for sample structure in genome-wide association studies. Nat Genet 42(4): 348–354.

Kang Y, Sakiroglu M, Krom N, Stanton-Geddes J, Wang M, Lee YC, Young ND, Udvardi M. 2015. Genome-wide association of drought-related and biomass traits with HapMap SNPs in Medicago truncatula. Plant Cell Environ. 38(10):1997–2011.

Kryvoruchko IS, Sinharoy S, Torres-Jerez I, Sosso D, Pislariu CI, Guan D, Murray J, Benedito VA, Frommer WB, Udvardi MK. 2016. MtSWEET11, a Nodule-Specific Sucrose Transporter of Medicago truncatula. Plant Physiol 171(1): 554–565.

Liang Y, Cao Y, Tanaka K, Thibivilliers S, Wan J, Choi J, Kang C, Qiu J, Stacey G. 2013. Nonlegumes respond to rhizobial Nod factors by suppressing the innate immune response. Science 341(6152): 1384–1387.

Liang Y, Tóth K, Cao Y, Tanaka K, Espinoza C, Stacey G. 2014. Lipochitooligosaccharide recognition: an ancient story. New Phytol 204(2): 289–296.

Limpens E, van Zeijl A, Geurts R. 2015. Lipochitooligosaccharides modulate plant host immunity to enable endosymbioses. Annu Rev Phytopathol 53: 311–334.

Maillet F, Poinsot V, André O, Puech-Pages V, Haouy A, Gueunier M, Cromer L, Giraudet D, Formey D, Niebel A, et al. 2011. Fungal lipochitooligosaccharide symbiotic signals in arbuscular mycorrhiza. Nature 469(7328): 58–63.

Nakagawa T, Kaku H, Shimoda Y, Sugiyama A, Shimamura M, Takanashi K, Yazaki K, Aoki T, Shibuya N, Kouchi H. 2011. From defense to symbiosis: limited alterations in the kinase domain of LysM receptor-like kinases are crucial for evolution of legume-Rhizobium symbiosis. Plant J. 65(2):169–80.

Ohsten Rasmussen M, Hogg B, Bono JJ, Samain E, Driguez H. 2004. New access to lipo-chitooligosaccharide nodulation factors. Org Biomol Chem 2(13): 1908–1910.

Olah B, Briere C, Bécard G, Dénarié J, Gough C. 2005. Nod factors and a diffusible factor from arbuscular mycorrhizal fungi stimulate lateral root formation in Medicago truncatula via the DMI1/DMI2 signalling pathway. Plant J 44(2): 195–207.

Pecrix Y, Staton SE, Sallet E, Lelandais-Brere C, Moreau S, Carrere S, Blein T, Jardinaud MF, Latrasse D, Zouine M, et al. 2018. Whole-genome landscape of Medicago truncatula symbiotic genes. Nature Plants 4(12): 1017–1025.

Rao S, Zhou Z, Miao P, Bi G, Hu M, Wu Y, Feng F, Zhang X, Zhou JM. 2018. Roles of Receptor-Like Cytoplasmic Kinase VII Members in Pattern-Triggered Immune Signaling. Plant Physiol 177(4): 1679–1690.

Rey T, Andre O, Nars A, Dumas B, Gough C, Bottin A, Jacquet C. 2019. Lipo-chitooligosaccharide signalling blocks a rapid pathogen-induced ROS burst without impeding immunity. New Phytologist 221(2): 743–749.

Rey T, Bonhomme M, Chatterjee A, Gavrin A, Toulotte J, Yang W, André O, Jacquet C, Schornack S. 2017. The Medicago truncatula GRAS protein RAD1 supports arbuscular mycorrhiza symbiosis and Phytophthora palmivora susceptibility. J Exp Bot 68(21-22): 5871–5881.

Ristova D, Giovannetti M, Metesch K, Busch W. 2018. Natural genetic variation shapes root system responses to phytohormones in Arabidopsis. Plant J 96(2): 468–481.

Roche P, Debellé F, Maillet F, Lerouge P, Faucher C, Truchet G, Dénarié J, Promé JC. 1991a. Molecular basis of symbiotic host specificity in Rhizobium meliloti: nodH and nodPQ genes encode the sulfation of lipo-oligosaccharide signals. Cell 67(6): 1131–1143.

Roche P, Lerouge P, Ponthus C, Prome JC. 1991b. Structural determination of bacterial nodulation factors involved in the Rhizobium meliloti-alfalfa symbiosis. J Biol Chem 266(17): 10933–10940.

Ronfort J, Bataillon T, Santoni S, Delalande M, David JL, Prosperi JM. 2006. Microsatellite diversity and broad scale geographic structure in a model legume: building a set of nested core collection for studying naturally occurring variation in Medicago truncatula. BMC Plant Biol 6: 28.

Rush T, Puech-Pagès V, Bascaules A, Jargeat P, Maillet F, Haouy A, Maës A, Carrera Carriel C, Khokhani D, Keller-Pearson M, et al. 2020. Lipo-chitooligosaccharides as regulatory signals of fungal growth and development. Nature Commun in press.

Samain E, Chazalet V, Geremia RA. 1999. Production of O-acetylated and sulfated chitooligosaccharides by recombinant Escherichia coli strains harboring different combinations of nod genes. J Biotechnol 72(1-2): 33–47.

Samain E, Drouillard S, Heyraud A, Driguez H, Geremia RA. 1997. Gram-scale synthesis of recombinant chitooligosaccharides in Escherichia coli. Carbohydr Res 302(1-2): 35–42.

Schiessl K, Lilley JLS, Lee T, Tamvakis I, Kohlen W, Bailey PC, Thomas A, Luptak J, Ramakrishnan K, Carpenter MD, et al. 2019. NODULE INCEPTION Recruits the Lateral Root Developmental Program for Symbiotic Nodule Organogenesis in Medicago truncatula. Curr Biol. 29(21):3657-3668.e5

Souleimanov A, Prithiviraj B, Smith DL. 2002. The major Nod factor of Bradyrhizobium japonicum promotes early growth of soybean and corn. J Exp Bot 53(376): 1929–1934.

Soyano T, Shimoda Y, Kawaguchi M, Hayashi M. 2019. A shared gene drives lateral root development and root nodule symbiosis pathways in Lotus. Science 366(6468): 1021–1023.

Stanton-Geddes J, Paape T, Epstein B, Briskine R, Yoder J, Mudge J, Bharti AK, Farmer AD, Zhou P, Denny R, et al. 2013. Candidate genes and genetic architecture of symbiotic and agronomic traits revealed by whole-genome, sequence-based association genetics in Medicago truncatula. PLoS One 8(5): e65688.

Stegmann M, Monaghan J, Smakowska-Luzan E, Rovenich H, Lehner A, Holton N, Belkhadir Y, Zipfel C. 2017. The receptor kinase FER is a RALF-regulated scaffold controlling plant immune signaling. Science 355(6322): 287–289.

Sulima AS, Zhukov VA, Afonin AA, Zhernakov AI, Tikhonovich IA, Lutova LA. 2017. Selection Signatures in the First Exon of Paralogous Receptor Kinase Genes from the Sym2 Region of the Pisum sativum L. Genome. Frontiers in Plant Science 8: 1957

Sulima AS, Zhukov VA, Kulaeva OA, Vasileva EN, Borisov AY, Tikhonovich IA. 2019. New sources of Sym2 A allele in the pea (Pisum sativum L.) carry the unique variant of candidate LysM-RLK gene LykX. PeerJ 7: e8070.

Sun J, Miller JB, Granqvist E, Wiley-Kalil A, Gobbato E, Maillet F, Cottaz S, Samain E, Venkateshwaran M, Fort S, et al. 2015. Activation of symbiosis signaling by arbuscular mycorrhizal fungi in legumes and rice. Plant Cell 27(3): 823–838.

Tanaka K, Cho SH, Lee H, Pham AQ, Batek JM, Cui S, Qiu J, Khan SM, Joshi T, Zhang ZJ, et al. 2015. Effect of lipo-chitooligosaccharide on early growth of C4 grass seedlings. J Exp Bot 66(19): 5727–5738.

Trujillo DI, Silverstein KA, Young ND. 2014. Genomic characterization of the LEED..PEEDs, a gene family unique to the medicago lineage. G3 (Bethesda) 4(10): 2003–2012.

Wang D, Couderc F, Tian CF, Gu W, Liu LX, Poinsot V. 2018. Conserved Composition of Nod Factors and Exopolysaccharides Produced by Different Phylogenetic Lineage. Front Microbiol 9: 2852.

Wang L, Wu N, Zhu Y, Song W, Zhao X, Li Y, Hu Y. 2015. The divergence and positive selection of the plant-specific BURP-containing protein family. Ecol Evol 5(22): 5394–5412.

Yamaguchi S. 2008. Gibberellin metabolism and its regulation. Annu Rev Plant Biol 59: 225–251.

Yoder JB, Stanton-Geddes J, Zhou P, Briskine R, Young ND, Tiffin P. 2014. Genomic signature of adaptation to climate in Medicago truncatula. Genetics 196(4): 1263–1275.

