## supplemental Figures for "Distinct genetic bases for plant root responses to lipo-chitooligosaccharide signal molecules from distinct microbial origins"

### « Fung - LCOs »

### Fucosylated or methyl-fucosylated (1:1 ratio)

### Nod-LCOs from *Sinorhizobium meliloti:*

### sulphated and acetylated

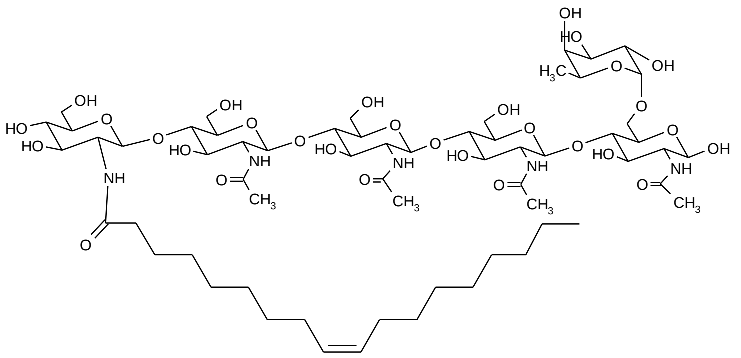

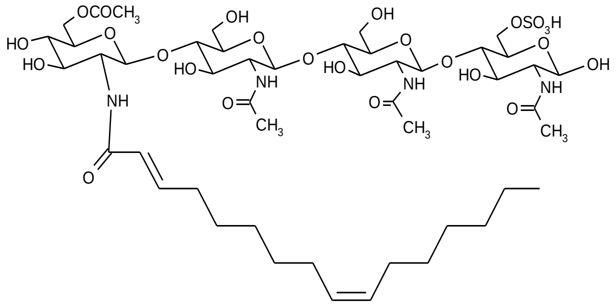

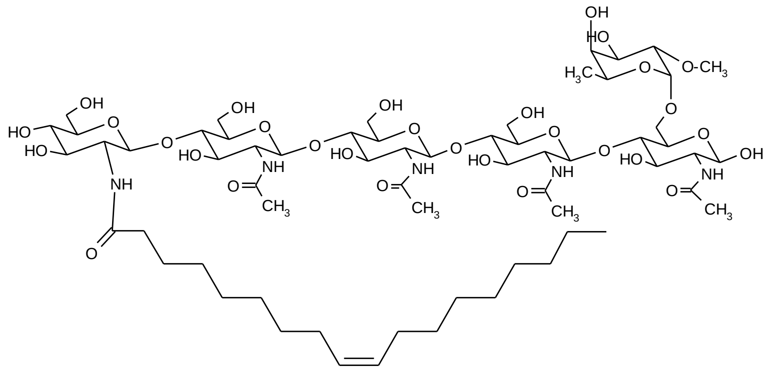

### « Myc - LCOs »

### Non sulfated or sulfated

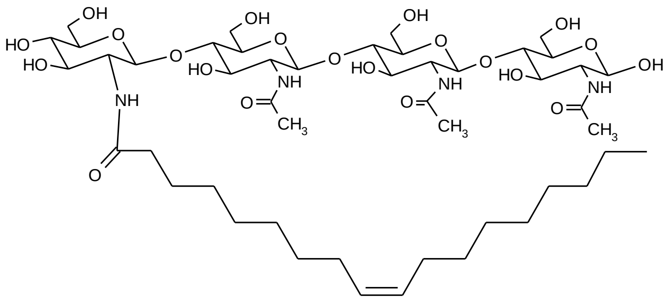

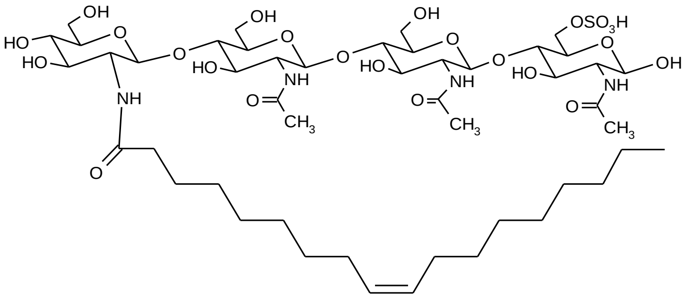

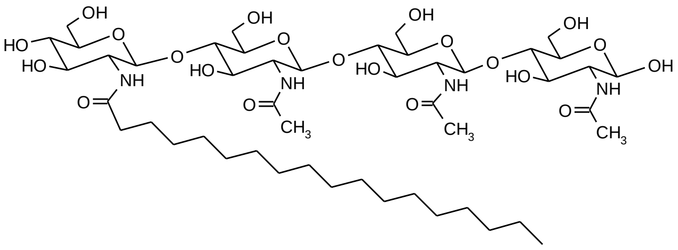

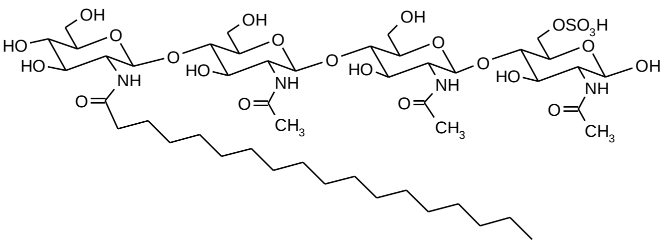

**Fig. S1**

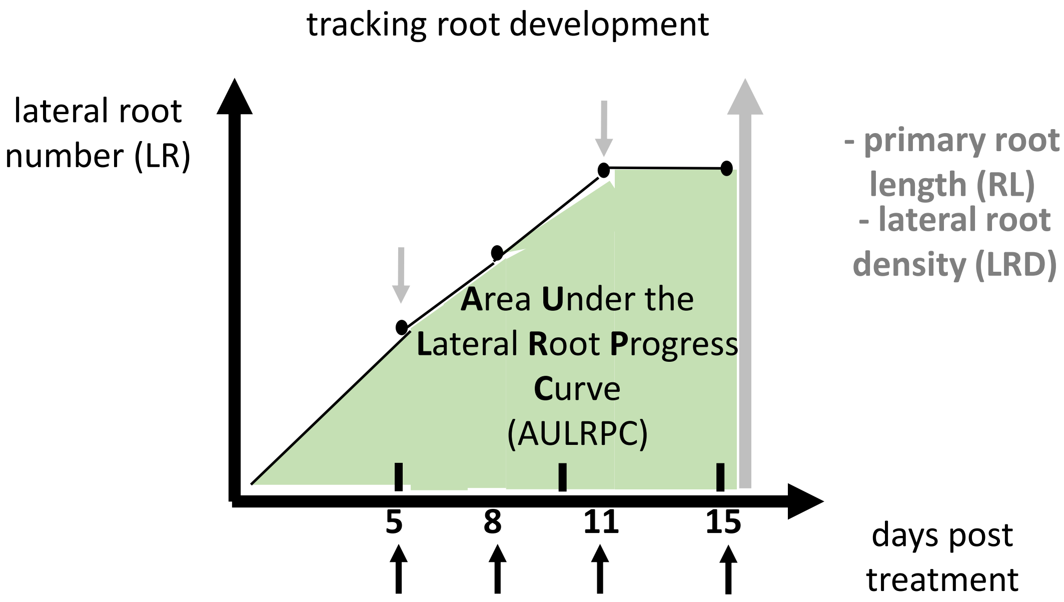
tracking root development
lateral root number (LR)
- primary root length (RL)
- lateral root density (LRD)
Area Under the Lateral Root Progress Curve (AULRPC)
5
8
11
15
days post treatment

Fig. S2
